# Multimodal neural correlates of cognitive control in the Human Connectome Project

**DOI:** 10.1101/124123

**Authors:** Dov B. Lerman-Sinkoff, Jing Sui, Srinivas Rachakonda, Sridhar Kandala, Vince D. Calhoun, Deanna M. Barch

## Abstract

Cognitive control is a construct that refers to the set of functions that enable decisionmaking and task performance through the representation of task states, goals, and rules. The neural correlates of cognitive control have been studied in humans using a wide variety of neuroimaging modalities, including structural MRI, resting-state fMRI, and task-based fMRI. The results from each of these modalities independently have implicated the involvement of a number of brain regions in cognitive control, including dorsal prefrontal cortex, and frontal parietal and cingulo-opercular brain networks. However, it is not clear how the results from a single modality relate to results in other modalities. Recent developments in multimodal image analysis methods provide an avenue for answering such questions and could yield more integrated models of the neural correlates of cognitive control. In this study, we used multiset canonical correlation analysis with joint independent component analysis (mCCA+jICA) to identify multimodal patterns of variation related to cognitive control. We used two independent cohorts of participants from the Human Connectome Project, each of which had data from four imaging modalities. We replicated the findings from the first cohort in the second cohort using both independent and predictive analyses. The independent analyses identified a component in each cohort that was highly similar to the other and significantly correlated with cognitive control performance. The replication by prediction analyses identified two independent components that were significantly correlated with cognitive control performance in the first cohort and significantly predictive of performance in the second cohort. These components identified positive relationships across the modalities in neural regions related to both dynamic and stable aspects of task control, including regions in both the frontal-parietal and cingulo-opercular networks, as well as regions hypothesized to be modulated by cognitive control signaling, such as visual cortex. Taken together, these results illustrate the potential utility of multi-modal analyses in identifying the neural correlates of cognitive control across different indicators of brain structure and function.

## Introduction

Cognitive control refers to the set of cognitive functions that are employed to encode and maintain task representations so as to regulate one’s thoughts and actions (Botvinick and Braver 2015). These functions are accomplished through the recruitment of neural systems that are also involved in supporting memory, perception, attention, action selection and inhibition, among other functions (Miller and Cohen 2001, Botvinick and Braver 2015). Together, these functions enable and regulate the decision-making processes that are omnipresent in life. Within the neuroimaging literature, several different imaging modalities have been used to study the neural underpinnings of cognitive control, including structural, functional, and resting state MRI. However, in much of the literature, a single neuroimaging modality is examined in a given study. This can make it difficult to understand how findings in different modalities relate to each other and to cognitive control. Thus, the goal of the present study was to use a data-driven multimodal analysis approach to study the neural correlates of cognitive control.

### Single Imaging Modality Studies

As noted above, much of the existing literature on the neural correlates of cognitive control have examined one imaging modality in a particular study. For example, a metaanalysis of 31 studies of cortical volume and 10 studies of cortical thickness in prefrontal cortex (PFC) revealed a moderate positive relationship between overall PFC volume and better cognitive control performance (Yuan and Raz 2014), with subregion analyses suggesting stronger relationships in lateral and medial PFC versus orbitofrontal cortex. Further, there was a significant relationship between PFC thickness and cognitive control, though there were not enough studies to examine the relationship between the thickness of subregions of PFC and cognitive control. Additional studies not included in this meta-analysis are consistent with these findings (Burzynska, Nagel et al. 2012, Tu, Chen et al. 2012), though the specificity of such relationships to PFC remains an open question.

Additionally, various forms of functional MRI (fMRI) have also been used to study cognitive control. While a full review of the task fMRI (tfMRI) literature is beyond the scope of this introduction (see (Niendam, Laird et al. 2012, Botvinick and Braver 2015, D’Esposito and Postle 2015), among others), meta-analytic evidence from this literature also strongly implicates prefrontal cortex areas as critical to cognitive control (Niendam, Laird et al. 2012). Drawing from 193 studies of cognitive control in healthy participants, Niendam and colleagues identified robust activation in lateral and medial prefrontal, dorsal anterior cingulate, and parietal cortex in response to a broad set of cognitive control paradigms. Further, they divided the studies into specific domains of cognitive control, which identified differential patterns of activation across these same areas as well as portions of the basal ganglia and cerebellum.

Resting state functional connectivity MRI (rsfcMRI) has also been used to study the neural correlates of cognitive control. For example, (Cole, Yarkoni et al. 2012) used global brain connectivity, a measure of a region’s connectivity with the rest of the brain, to identify a region in lateral prefrontal cortex wherein resting activity was highly correlated with fluid intelligence, an index related to cognitive control. (Seeley, Menon et al. 2007) used an ROI and ICA based approach to rsfcMRI and identified clusters in bilateral intraparietal sulcus that positively correlated with better cognitive control. Further, recently developed methods in dynamic rsfcMRI (Calhoun, Miller et al. 2014) have identified specific modes of neural resting-state connectivity and that inter-individual differences in the tendencies to use particular modes of connectivity were related to cognitive control. Specifically, modes which showed strong modular networks and anticorrelated relationships from visual and somatosensory areas to cerebellar regions, were significantly correlated with improved performance on several executive tasks including measures of cognitive flexibility, processing speed, and working memory but not with fluid intelligence or inhibition and attention (Nomi, Vij et al. 2016).

As reviewed above, analyses of structural, functional, and connectivity relationships to cognitive control have often identified overlapping regions. For example, both the structural and functional activation meta-analyses point to lateral and medial regions of prefrontal cortex, as have some of the functional connectivity studies. However, what is not clear is whether these are the same regions of prefrontal cortex across modalities or studies, and whether they correlate across individuals. Further, how do patterns in large-scale network organization from rsfcMRI data in and between those regions relate to measures of cortical thickness and functional activation? How do these patterns across different imaging modalities relate with behavior? These questions are difficult to answer with single modality studies, and their answers could provide broader insights into neural functions.

### Examining Multiple Modalities

Given the complimentary strengths and weaknesses associated with each modality (Biessmann, Plis et al. 2011), many studies collect several different imaging modalities in the same individual, often in the same scanning session. However, many investigators choose to analyze these different imaging modalities using independent analysis pathways (Groves, Beckmann et al. 2011). With such an approach, the integration of findings occurs post-hoc using approaches such as correlation between measures or visual inspection and description (Groves, Smith et al. 2012, Calhoun and Sui 2016). For example, (Westlye, Walhovd et al. 2009) correlated the results of independently processed DTI data with EEG data from a flanker task which identified a significant relationship between the two modalities in the posterior left cingulum. Similarly, (Harms, Wang et al. 2013) used a post-hoc correlation based approach and identified a relationship between volume of the superior and middle frontal gyri and working memory related activity in the intraparietal sulcus and a relationship between hippocampal volume and working memory related activity in the dorsal anterior cingulate and left inferior frontal gyrus (Harms, Wang et al. 2013).

While such correlational approaches are important and have yielded informative results, they represent a univariate approach to a multivariate problem (Calhoun and Sui 2016). This can generate a unique set of findings within a given modality with relatively little guidance as to how the results fit together across modalities (Sui, Adali et al. 2012, Pearlson, Liu et al. 2015, Calhoun and Sui 2016). As shown in (Calhoun and Sui 2016), data from (Plis, Weisend et al. 2011) were used to perform independent analyses in fMRI and MEG data that were collected from the same set of subjects performing the same task. These data were used to generate network graph representations for both modalities independently and resulted in graphs with highly dissimilar structures and properties. In contrast, combined multimodal analysis using the same data led to brain networks in the individual modalities that were highly spatially correlated. While further data are needed to determine whether one type of analysis approach versus the other is better related to external validators, the findings do suggest the univariate approach to multimodal data analysis does not always identify coherent patterns across modalities.

### Multimodal Fusion Analysis Approaches

To address this, recent methodological advances have provided a new set of analysis tools aimed towards solving the difficulties in adjudicating between dissimilar results generated by analyzing multiple modalities in separate pathways (Michael, Baum et al. 2010, Biessmann, Plis et al. 2011, Groves, Beckmann et al. 2011, Sui, Adali et al. 2012, Calhoun and Sui 2016). These methods enable analysis of multiple imaging modalities in a single analysis, which allows for simultaneous study of the brain at multiple levels of analysis and capitalizes on the complimentary strengths across modalities (Biessmann, Plis et al. 2011). Further, these approaches are able to identify joint variance structures that help us understand the shared patterns contained within the different modalities of data and can present a richer understanding of the neural constructs under examination (Sui, He et al. 2012).

One such method is multiset canonical correlation analysis with joint independent component analysis (mCCA+jICA) (Sui, Pearlson et al. 2011, Sui, He et al. 2012, Sui, He et al. 2013). This method simultaneously decomposes multiple modalities of data and identifies a set of hidden sources of variance that are linked across modalities and jointly contribute to the variation seen in the data. The combination of these two analysis methods, mCCA (Li, Adali et al. 2009) and jICA (Calhoun, Adali et al. 2006), overcomes the limitations of the individual methods (see (Sui, Adali et al. 2012) for review) and provides a mathematical framework that enables the identification of strong and weak linkages across modalities as well as the identification of modality-unique features in the data (Sui, He et al. 2012). Together, these methods identify maximally independent, cross-modality linked sources of variance (independent components [ICs]) in the data as well as subject-specific weights upon the group-level ICs. These weights can then be used in post-hoc analyses of individual differences allowing for the determination of multimodal brain and behavior relationships.

### Replication

mCCA+jICA and other multimodal analysis methods are powerful, data-driven methods designed to detect complex patterns hidden within data (Calhoun and Sui 2016). However, this power comes with a risk of overfitting results to a particular sample such that the results may not generalize to other participant samples. Indeed, concerns for replicability are growing in the psychological and neuroimaging literature (Barch and Yarkoni 2013, Open_Science_Collaboration 2015) as well as the broader scientific literature (Baker 2016). Given this risk, it is becoming increasingly important to design data-driven studies with replication in mind. One avenue for addressing this is to design analyses around extant datasets that have large numbers of subjects and rigorous quality control processes, such as the Human Connectome Project (Van Essen, Smith et al. 2013), or through open sharing of data on platforms such as the OASIS database (www.oasis-brains.org), the COINS platform (Landis, Courtney et al. 2016) (http://coins.mrn.org), or the OpenfMRI project (Poldrack, Barch et al. 2013) (https://openfmri.org/). Further, given the large numbers of subjects in some of these databases, studies can be designed with built-in replication through a variety of methods. These methods range from the straightforward, such as splitting data into two cohorts with independent analyses and post-hoc comparisons, to predictive analyses, where the results from one cohort are used to predict another cohort, to more complicated methods from the machine learning literature such as k-fold cross-validation.

### Current Study

Thus, the goal of the present paper was to use mCCA+jICA to examine the multi-modal neural correlates of cognitive control in a healthy community sample using data from the Human Connectome Project (HCP). We selected two cohorts of participants from the HCP (n=194 and n=149), each of which contained complete behavioral and imaging data in our domains of interest. These cohorts were selected to enable two types of replication analyses. First, we analyzed Cohort 1 by extracting imaging features for participants from four imaging modalities (sMRI, rsfcMRI, and two tfMRI tasks) and applying mCCA+jICA to these imaging features. Analyses yielded a set of group-level modality-linked independent sources of variance as well as individual subject weightings on these sources. These weights were then correlated with a composite behavioral metric of cognitive control. We performed the first replication analysis by using the group-level results from the first cohort of subjects to predict the results of Cohort 2. We then performed the second replication analysis by independently analyzing the second cohort of subjects with mCCA+jICA. We used a similarity algorithm to match visual patterns across the two cohorts, and then correlated the second cohort’s independently derived subject-specific weights with the composite behavioral metric of cognitive control.

## Methods

### Participants

The present study drew participants from the HCP database (Van Essen, Smith et al. 2013). Briefly, participants recruited to participate in the HCP were a healthy community sample approximating the demographics of the United States. Participants were between the ages of 22-35 with no documented history of mental illness, neurological disorder, or physical illness with known impact upon brain functioning. Additionally, participants had no contraindications to the MRI environment. We selected a cohort of participants from the HCP such that there were no related participants within the cohort due to concerns of the heritability of neural features (Glahn, Winkler et al. 2010). This yielded n=194 participants in this cohort (cohort 1). A second cohort of participants (n=149) was selected for use in two types of replication analyses (cohort2). There were no differences across cohorts in age, gender, or years of education (table S1). ID numbers for the participants used in each of these cohorts are available in the supplement. Due to technical issues, one subject contributed imaging data but did not contribute behavioral performance data. Thus, for Cohort 1, n=194 subjects contributed imaging data and n=193 subjects were used for statistical analyses with behavior.

### Behavioral assessment

For each participant, we computed a composite measure of cognitive control as the summed z-score of four behavioral metrics, with each of these tasks described in detail in prior work (Barch, Burgess et al. 2013). These measures were: (1) N-back working memory task – accuracy in the 2-back condition. In this task, participants were presented a mixture of four different stimulus types (faces, places, tools, and body parts) and were asked to respond when the displayed stimulus was the same as the stimulus displayed two stimuli prior. Twenty percent of presented stimuli were targets and 20-30% were lures (target stimuli in 1-back or 3-back conditions) to ensure that participants used an active memory approach rather than a passive familiarity approach. (2) Relational processing task – accuracy in the relational condition. This task was a modified version of the task used in (Smith, Keramatian et al. 2007). Participants were presented with two pairs of stimuli (6 possible shapes with 6 possible textures) with one pair on the top of the screen and the other on the bottom. Participants were instructed to determine whether the top pair shared the same shape or texture, and then, whether the bottom pair varied along the same dimension. (3) Flanker task scaled score from the NIH Toolbox (Gershon, Wagster et al. 2013, Hodes, Insel et al. 2013). Participants were presented with collinear directional arrows and instructed to attend to the central arrow. Participants were asked to respond by indicating whether the central arrow pointed left or right, while ignoring the direction of the flanking arrows. Flanker scores were a normed combination of accuracy and reaction time. (4) Penn Progressive Matrices – total number of correct responses. This was a shortened version of the classic Raven’s progressive matrices test for fluid intelligence (form A (Bilker, Hansen et al. 2012)). Participants were presented with a texture with a section removed and asked to determine which one of six possible options would fit the pattern of the texture.

We assessed the internal consistency of our composite measure of cognitive control performance using SPSS 23 (IBM, Armonk, NY). The individual z-scored behavioral performance metrics from both cohorts were pooled and Cronbach’s alpha was found to be 0.64 using all four metrics. When working memory task performance, relational task performance, or progressive matrices task performance were deleted from the composite, Cronbach’s alpha decreased to 0.44, 0.54, and 0.57 respectively. In contrast, Cronbach’s alpha increased marginally to 0.7 with the removal of the flanker task performance. However, given that the increase in internal consistency was marginal, all four metrics were retained in the final composite measure of cognitive control.

### Image collection and feature selection

All imaging data were collected and pre-processed as part of the HCP. Briefly, all scanning was performed on a 3T customized Siemens “Connectom” Skyra scanner with a 32-channel head coil and 100 mT/m gradient coils. T1 and T2 images were acquired at 0.7 mm isotropic resolution. BOLD contrast images were acquired using a gradient-echo echo-planar 8X multiband accelerated sequence with 2mm^3^ isotropic voxels (TR=720ms). Resting state data were collected over two days in four 15-minute sessions with eyes open and crosshair fixation (Van Essen, Smith et al. 2013). Task data were collected over two days. The working memory (2-back) task duration was 602 seconds and the relational processing task duration was 352 seconds (combined L->R and R->L phase-encoding scans) (Barch, Burgess et al. 2013).

Participant’s structural scans were collected and processed through the HCP’s minimal preprocessing pipelines as described in (Glasser, Sotiropoulos et al. 2013). Briefly, T1 and T2 weighted images were processed through three sequential HCP structural-image pipelines. The initial pipeline performed the following: corrected gradient nonlinearity-induced distortions; aligned subject native-space scans to MNI coordinate space; removed readout-distortions; corrected intensity inhomogeneities; and then aligned native-space data to the MNI atlas. Next, the second pipeline processed participant data through a customized version of Freesurfer to generate subject-specific brain segmentations and parcellations that take advantage of the HCP’s high-resolution structural data. Finally, the third and final pipeline converted Freesurfer output files into NIFTI, CIFTI, and GIFTI formats, as well as registered data to several different surface meshes, including the 32k surface mesh that was used as the standard space for all downstream analyses in the present report. The present study used Freesurfer-determined measures of cortical thickness at every vertex in the 32k surface mesh as the sMRI measure.

Participants task (tfMRI) and resting state (rsfcMRI) scans were collected (Ugurbil, Xu et al. 2013) and processed identically through the HCP’s pipelines to generate data aligned to the 32k surface mesh (Glasser, Sotiropoulos et al. 2013). Briefly, for each task, task runs were acquired in two phase-encoding directions (L->R, R->L) and resting state data were acquired in four runs (2 of each phase-encoding direction). All functional data were then first processed through a volume minimal-preprocessing stream (Glasser, Sotiropoulos et al. 2013) configured to perform gradient unwarping, motion correction, EPI field distortion correction, registration to T1w data, registration into MNI space, and intensity normalization. The cortical ribbon was then projected to the surface and registered with the structural meshes from the structural pipelines into a standard “grayordinates” surface space while also including a set of volumetric data for subcortical and cerebellar regions. Surface- and volume-based smoothing algorithms were applied to bring total smoothing to 4mm FWHM. The minimal preprocessing pipelines for rsfcMRI data ended here (see below for further rsfcMRI processing details). tfMRI data were analyzed using FSL to generate subject-specific spatial map COPEs corresponding to the desired task condition. In the present study, we used the working memory task: 2-back condition contrast; and the relational processing task: relational condition contrast (tasks described above in *behavioral*).

Resting state data from the HCP were further processed in-house to generate correlation matrices for each subject. Participants’ resting state data were demeaned and detrended within each run. Twenty-four motion regressors (6 motion parameters, their derivatives, and squares), along with the unique noise components from MELODIC identified by FIX (Griffanti, Salimi-Khorshidi et al. 2014, Salimi-Khorshidi, Douaud et al. 2014), and the mean grayordinates timeseries (global signal) and its first derivative were removed in a single regression. Data were then processed through a highpass filter (cutoff = 0.009 Hz). Data were demeaned and detrended, and no additional spatial smoothing was applied (Burgess, Kandala et al. 2016). Runs were then concatenated, and mean timeseries were extracted from a known cortical surface parcellation scheme (Gordon, Laumann et al. 2016), with the addition of several parcels from the cerebellum (Culbreth, Kandala et al. 2016) and subcortical regions defined by Freesurfer. Functional connectivity matrices were then computed as the Pearson correlation between all parcels.

### mCCA+jICA multimodal imaging analysis

The four data features described above (cortical thickness, resting state functional connectivity correlations, and task COPEs) were used as the imaging measures of interest for mCCA+jICA. Multiset canonical correlation analysis + joint independent component analysis (mCCA+jICA) is a blind source separation method that simultaneously decomposes multiple modalities of data to reveal independent latent sources of variance in the data. It is a flexible analysis method capable of identifying the modality-unique and cross-modality patterns of variance within the data (Sui, Adali et al. 2012). In the initial step, mCCA, an extension of traditional canonical correlation analysis, projects the data into a space that links the imaging modalities to maximize inter-subject covariation. Following that, the components from mCCA are further decomposed in a joint ICA framework in order to identify maximally independent sources of variance.

All feature extraction, analyses, post-processing, and visualization were performed in MATLAB (The Mathworks, Natick, MA) R2012b and R2015a using custom-written code and an in-house modified version of the FIT toolbox (base version 2.0.c) (publicly available at http://mialab.mrn.org/software/fit/). MATLAB-native visualization tools for surface data as well as an interactive resting state data viewer are available on github (code to be released upon acceptance of manuscript). For each subject, data files corresponding to cortical thickness, rsfcMRI matrices, and the two tfMRI z-scored COPEs from FSL were loaded into MATLAB, linearized into vectors, and stacked into four matrices (one matrix per imaging modality) such that matrices were of size N subjects-by-number of features (voxels and/or vertices) in the respective modality. Dimensionality of the data matrices was then reduced using a singular value decomposition (see *parameter sweep*, in supplemental methods) that maintained high percentages of accounted variance (see *Parameter Sweep results*, in supplement) and then analyzed using mCCA.

mCCA (Correa, Li et al. 2008, Li, Adali et al. 2009) is a multimodal extension of canonical correlation analysis. Within the broader framework of mCCA+jICA, the goal of the mCCA step is to align the data such that it simplifies the correlational structure across the modalities and maximizes inter-subject covariation (Sui, Pearlson et al. 2015). In doing so, mCCA decomposes each modality into a set of mixing profiles (the subject loading parameters) and the corresponding components (spatial maps). The mixing profiles (loading parameters) contain a set of weights that describe how much of a given component is required to reconstruct an individual subject’s source data. The components contain spatial maps that represent how strongly a given voxel/vertex is weighted relative to the other voxels/vertices in the spatial map, and can be interpreted similar to standard fMRI spatial maps. Through an iterative multi-step process, mCCA maximizes a sum of squares of correlations cost function such that the corresponding canonical variants (CV) across the four modalities are maximally correlated. That is, CV1 for sMRI data is maximally correlated with CV1 for rsfcMRI, CV1 for relational tfMRI, and CV1 for 2-back tfMRI, but not with CVs 2 through M (where M is the final number of components). In linking CVs, this process also links the corresponding components (spatial maps) across modalities. However, while the components are linked, mCCA may fail to achieve fully separated sources when applied to neuroimaging data due to underlying noise and dependencies in the data (Correa, Adali et al. 2010, Sui, Adali et al. 2010, Sui, Pearlson et al. 2011).

To overcome this incomplete separation of sources, the data were further decomposed into maximally spatially independent sources of variance through the application of joint Independent Component Analysis (jICA). jICA is a extension of traditional ICA methods (Calhoun, Adali et al. 2006) that identifies latent sources of variance in the multimodal data. The set of component matrices (not CVs) from mCCA were joined into a single data matrix by concatenating along the feature (vertex/voxel) dimension, resulting in a component-by-feature matrix. jICA analyses were repeated 100 times with random initial conditions using the Infomax ICA algorithm (Bell and Sejnowski 1995) within the ICASSO framework (Himberg and Hyvärinen 2003, Himberg, Hyvärinen et al. 2004) to ensure stability and reproducibility of jICA analyses. Results of the 100 jICA analyses were grouped by component number (e.g.: components 1 through M), and the individual component estimate within a group that was most similar to all other component estimates within its group was selected as the component for further analysis.

Similarly to mCCA, jICA generates a set of matrices of subject-specific weights (N matrices, where N is number of modalities each of size subjects-by-number of components) as well as a matrix of components (of size components-by-features). The component matrices were interpreted by unstacking the matrices into the four individual modality matrices then and used to create standard CIFTI files for visualization with an in-house developed viewing tool. The subject-specific weights were extracted and used for statistical analyses with behavior.

### Statistical analyses

As described above, mCCA+jICA returns a set of subject-specific weights for each component. These weights were extracted and imported into SPSS 23 (IBM, Armonk, NY) for correlation analyses. These correlation analyses were performed only after all other analyses were completed (parameter determination via sweep [see supplementary methods],

mCCA+jICA analysis, and component visualization) in order to prevent bias in our selection of mCCA+jICA parameters. Weights for each modality within each component were independently correlated with our composite measure of cognitive control (see *behavioral assessment* above) and tested for significance using an FDR corrected two-tailed alpha = 0.05. FDR correction was performed in MATLAB using publicly available tools (http://www.mathworks.com/matlabcentral/fileexchange/27418).

### Replication by prediction

As noted above, we performed two different replication analyses. We wanted to assess whether the identified ICs had predictive power when applied to another cohort. This was assessed by using the ICs identified from Cohort 1 (fig. 1B) to decompose the source imaging data from Cohort 2 (fig. 1D). This process generated a set of subject-specific weights for Cohort 2 that corresponded to the extent that a given subject’s data from Cohort 2 could be represented by the components determined from Cohort 1 (fig. 1E). This was accomplished by multiplying Cohort 2’s source data by the pseudoinverse of Cohort 1’s ICs. These generated weights for Cohort 2 were then correlated with behavior as was described above.

**Figure 1.**
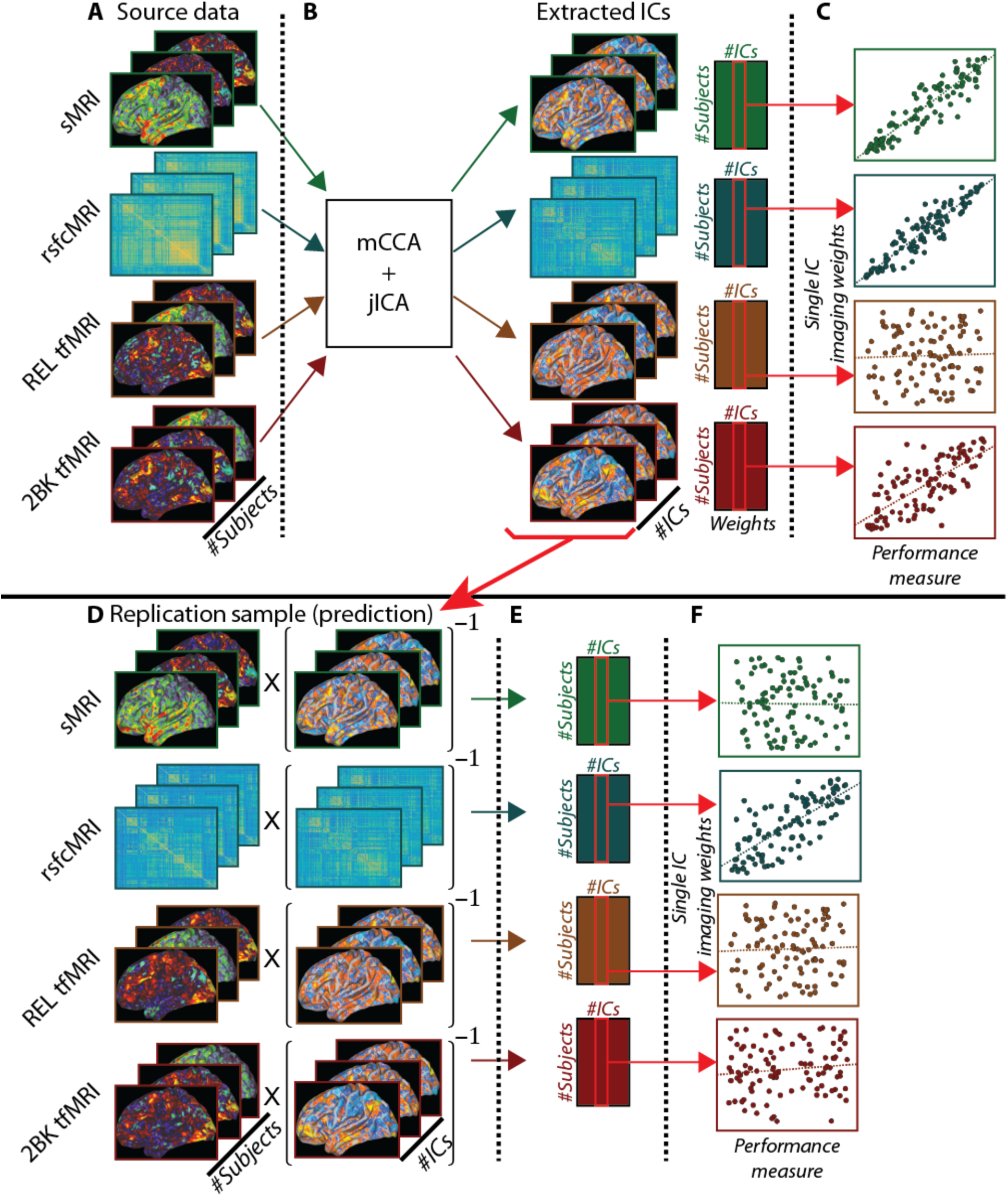
Schematic of analyses for independent and replication by prediction analyses. <u>Figure Caption</u>: Figure 1 is a schematic representation of the processing steps used to perform the present analyses. **Top:** analysis schematic for independent analyses in both cohorts. (A) Source imaging data. (B) mCCA+jICA analysis generated a set of group-level independent components (ICs) and subject-specific weights upon those ICs. (C) Subject specific weights from mCCA+jICA were used in correlation analyses with the performance measure. **Bottom:** analysis schematic for replication by prediction, using Cohort 1 to predict Cohort 2. (D) Source imaging data for Cohort 2 was multiplied by the pseudoinverse of the ICs from Cohort 1 (step 1B). (E) Subject-specific weights for Cohort 2 upon the imaging data (ICs) from Cohort 1. (F) Derived weights for Cohort 2 were correlated the performance measure.

### Replication by independent analysis

We additionally wanted to determine whether an independent application of mCCA+jICA to Cohort 2 would yield similar results to those found independently in Cohort 1. This analysis was performed in accordance with our model parameter selection criteria (see *supplementary methods*) in order to match the amount of accounted variance and across cohorts in the initial dimensionality reduction step. Given that the results of mCCA+jICA are data-driven and thus not identical for different analyses, we implemented a semi-automated cross-cohort component-matching algorithm to aid visual comparison of the ICs across the two cohorts. This algorithm used an eta^2^ similarity function (Cohen, Fair et al. 2008) that measured the amount of variance in one component that was accounted for by another component. This measure varied from 0 (complete dissimilarity) to 1 (identical components) and, unlike Pearson correlation, accounted for magnitude of differences between components as well as the covariance.

We first computed a matrix of eta^2^ values comparing the absolute value of each component in cohort 1 to each component in cohort 2. Absolute valued components were used in the eta^2^ computations instead of signed components because the mCCA+jICA model sometimes flips the sign of the components. This sign flipping is based upon a convention that treats the largest value in the component as positive even though it is also mathematically valid to leave the sign unflipped. We used a simple maximization of similarity algorithm that chose (without replacement) the highest value of similarity in the matrix and iterated through until all components were matched. The matched pairs of ICs were then visually examined and subject-specific weights correlated with the cognitive control behavioral metric.

## Results

### Behavioral data

As described in methods, all subjects performed four behavioral tasks that index cognitive control and were summed to create a composite. The distributions of the cognitive control composite values were compared across cohorts using a two-sample Kolmogorov-Smirnov test which showed no differences in the distribution (p>0.99) (table S2). Further, there were no significant differences between replication cohorts on the individual metrics of cognitive control (table S2) and histograms of the individual metrics and composite scores are available in the supplement (figs. S3 and S4).

### Decomposition results and images

We used mCCA+jICA to decompose imaging data for Cohort 1. This identified nine independent components (ICs) based upon the results of our parameter sweep (see supplementary methods and results). Each of these components contained four linked sources of variance: three spatial maps (corresponding to sMRI and the two tfMRI modalities) and a symmetric correlation matrix (corresponding to the rsfcMRI modality). For the three spatial maps, the value at each vertex/voxel in the spatial map corresponded to a weighting of how important that vertex/voxel was to that component, relative to all other vertices/voxels in that map. For rsfcMRI, each element in the correlation matrix corresponded to a weighting of how important that pairwise correlation between parcels was to that component, relative to all other pairwise correlations in the matrix.

### Correlations with behavior

In addition to generating a set of ICs, mCCA+jICA also generated a set of subject-specific weights (1 weight per-subject, per-IC, per-modality; 36 weights per participant) that describe the extent to which a given IC comprises the participant’s original data (less the variance lost in the dimensionality reduction). These weights were correlated with the composite measure of cognitive control with FDR used to correct for multiple comparisons. This revealed that all four imaging modalities strongly and significantly correlated with cognitive control performance after FDR correction for only one of the 9 ICs described above, C1-IC2 (Cohort 1 – Independent Component 2) (tables 1, S4). Examination of the corresponding scatterplots (fig. 3) showed that these correlations were not driven by outliers in the data.

**Table 1:**
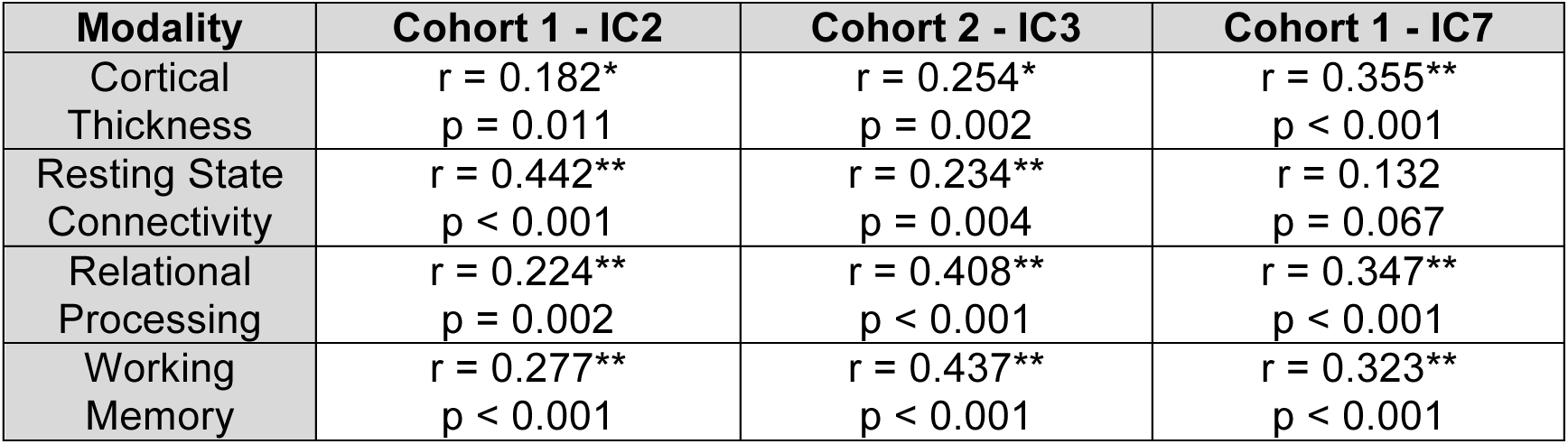
Correlations between subject-specific independent component weights and cognitive control composite behavioral metric for each cohort. <u>Figure Caption</u>: Correlations between subject-specific weights upon the independent components (IC) and the composite cognitive control behavioral metric revealed that Cohort 1’s IC2 and Cohort 2’s IC3 significantly correlated with behavior for all four imaging modalities, even after FDR correction. Cohort l’s IC7 was significantly correlated with behavior for three of the four imaging modalities. The ICs and correlation values presented here were generated in independent analyses of the cohorts. We did not identify an analog of C1-IC7 in Cohort 2. p-values in the table are original, unmodified values, all of which met or exceeded the critical p-value as determined by FDR (Cohort 1 p = 0.011; Cohort 2 p = 0.022). Correlation results for the other ICs are in the supplement. *p < 0.05, **p < 0.01, uncorrected.

| Modality | Cohort 1 - IC2 | Cohort 2 - IC3 | Cohort 1 - IC7 |
| --- | --- | --- | --- |
| Cortical Thickness | $r = 0.182^*$<br>$p = 0.011$ | $r = 0.254^*$<br>$p = 0.002$ | $r = 0.355^{**}$<br>$p < 0.001$ |
| Resting State Connectivity | $r = 0.442^{**}$<br>$p < 0.001$ | $r = 0.234^{**}$<br>$p = 0.004$ | $r = 0.132$<br>$p = 0.067$ |
| Relational Processing | $r = 0.224^{**}$<br>$p = 0.002$ | $r = 0.408^{**}$<br>$p < 0.001$ | $r = 0.347^{**}$<br>$p < 0.001$ |
| Working Memory | $r = 0.277^{**}$<br>$p < 0.001$ | $r = 0.437^{**}$<br>$p < 0.001$ | $r = 0.323^{**}$<br>$p < 0.001$ |

**Figure 2.**
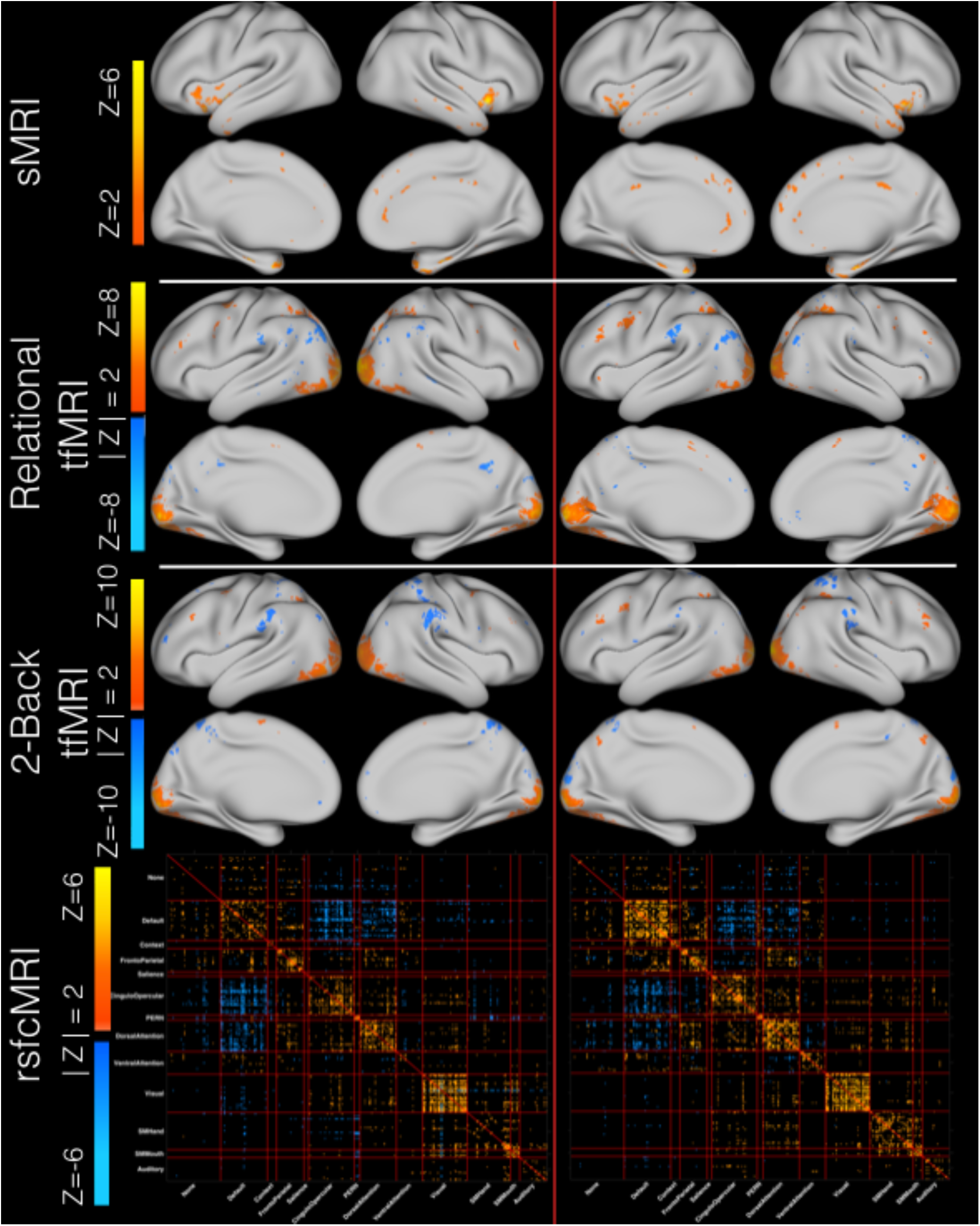
Imaging results for the similarity matched pair C1-IC2 and C2-IC3 for all four imaging modalities. <u>Figure Caption</u>: Figure 2 exhibits the spatial maps and correlation matrices that comprise Cohort 1’s IC2 (left column) and Cohort 2’s IC3 (right column). Modalities are the same across each row. All data are shown thresholded at |Z| > 2 with the exception of sMRI which is shown at Z > 2. For sMRI, the non-z-scored spatial maps were fully positive (Fig. S5). When Z-scored, the distribution was shifted to zero mean and thus only vertices where the signed Z-score value exceeded positive two are displayed, as the negatively valued Z-scored vertices represent those vertices that had the smallest magnitudes. Each modality is scaled independently to the minimum and maximum Z-value within a given modality for both cohorts in order to illustrate the strongest contributing vertices \ correlations within a given modality. Larger comparison images and the corresponding overlap masks are available in the supplement (Figs S5-12).

**Figure 3:**
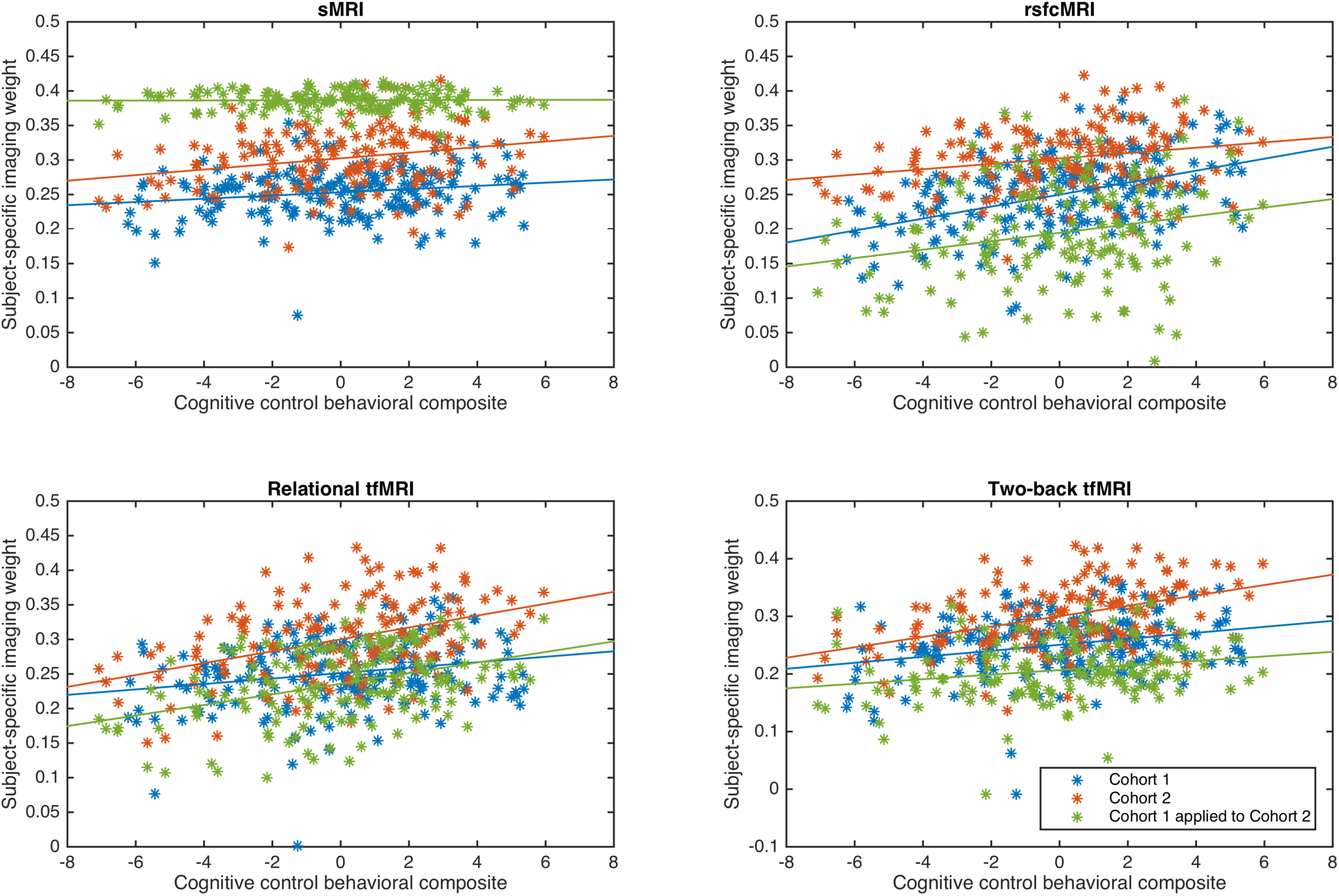
Scatter plots of cognitive control composite measure and subject-specific imaging weights for Cohort 1 independent component 2, Cohort 2 independent component 3, and the replication by prediction analysis for Cohort 1 independent component 2 predicting Cohort 2.

The mCCA+jICA model returns ICs with a value at every feature (i.e.: every vertex or pairwise correlation) in the map for each of the four modalities. To better understand these patterns, we converted the values in these maps to Z-scores and only displayed those vertices or pairwise correlations that exceeded a threshold of |Z| > 2 (Sui, Pearlson et al. 2015). Thus, the spatial maps and correlation matrices presented in figures 2, 4, S5-S15 represent those vertices or correlations that were strongest relative to all other vertices or correlations within their modality and IC. Further, there were minimal subcortical and cerebellar voxels exceeding the |Z| > 2 threshold for both tfMRI modalities; data at a threshold of |Z| > 1 are available for reference in the supplement (figs. S11, S12) (see *Limitations*).

For sMRI data, the strongest contributing areas in C1-IC2 were located predominantly in the bilateral insula, temporal poles, anterior middle temporal gyrus, right rostral anterior cingulate, right posterior cingulate, right isthmus of the cingulate, medial superior frontal cortex, and superior and inferior temporal gyri (fig. 2, S6). Thus, for sMRI data, greater cortical thickness in these areas was correlated with better cognitive control performance.

For relational tfMRI data (fig. 2, S7), the strongest contributing areas were located bilaterally in visual cortex, superior and inferior parietal cortex, inferior temporal cortex, left supramarginal gryus, left precentral sulcus, right rostral middle frontal cortex, bilateral superior precentral sulcus and gyrus, and inferiotemporal and fusiform gyri. This pattern suggests that for relational tfMRI, greater positive contributions in visual and superior parietal areas and greater negative contributions in the left inferior parietal areas were correlated with better cognitive control performance.

For 2-back tfMRI data (fig. 2, S8), the strongest positive contributing areas were in bilateral visual cortex and a small cluster in the left middlefrontal gyrus. Additionally, the strongest contributing negative clusters were in the bilateral supramarginal gyrus, and right superior parietal gyrus. This pattern suggests that for the 2-back tfMRI, greater positive contributions in visual areas and greater negative contributions in right parietal areas were correlated with better cognitive control performance.

For rsfcMRI data (fig. 2, S9, S10), the predominance of strongest positively correlated contributing connections were located along the diagonal of the matrix. This pattern suggests that better cognitive control performance is associated with a modular network structure wherein individual networks are more tightly connected to themselves than to other networks. Further, the matrix also contained strongly contributing anticorrelated connections between the default mode network (DMN) and task positive networks including the salience, cingulo-opercular, and dorsal attention networks. This pattern again suggests that better cognitive control was associated with stronger anti-correlations between the DMN and task positive networks. The rsfcMRI data exhibited robust subcortical and cerebellar correlations within those regions as well as with cortical parcels (fig. S10). Here too, there were concentrations of data along the diagonal (fig. S10 – sections A through R) suggesting that a more modular network structure was associated with better cognitive control. Further, there were strong within-network connections between the cortical and cerebellar structures for the DMN, PERN, fronto-parietal, and cingulo-opercular networks.

We further examined correlations between the subject-specific IC weights and each of the four individual measures of cognitive control that comprise the composite measure (table 2). Relational processing accuracy and number of correct responses on the progressive matrices task were significantly or trend-level correlated with all four imaging modalities. Accuracy on the two-back working memory task was significantly correlated with rsfcMRI data and both tfMRI modalities. The flanker task was only significantly correlated with rsfcMRI data.

**Table 2.** Individual behavioral task performance correlations with subject specific weights on C1-IC2 and C2-IC3. <u>Figure Caption</u>: Examination of individual behavioral metrics for Cohort 1 (top) revealed that the relational processing accuracy, progressive matrices task, and 2-back working memory task were correlated with all four modalities at significant or trend-level p-values. In contrast, the flanker task was significantly correlated with only resting state MRI. Examination of individual behavioral metrics for Cohort 2 (bottom) revealed that the relational processing accuracy and progressive matrices tasks were correlated with all four modalities at significant or trend-level p-values. While less significantly correlated, both 2-back working memory task accuracy and the flanker task correlated at trend-level or significant p-values for a subset of the modalities. *p < 0.05, **p < 0.01, uncorrected.

| **Cohort 1 - IC2** | **Cortical thickness** | **Resting state** | **Relational tfMRI** | **2-Back tfMRI** |
| --- | --- | --- | --- | --- |
| **Flanker task** | $r = 0.098$<br>$p = 0.174$ | $r = 0.262^{**}$<br>$p < 0.001$ | $r = -0.012$<br>$p = 0.872$ | $r = 0.047$<br>$p = 0.514$ |
| **2-Back accuracy** | $r = 0.130$<br>$p = 0.073$ | $r = 0.371^{**}$<br>$p < 0.001$ | $r = 0.193^{**}$<br>$p = 0.007$ | $r = 0.241^{**}$<br>$p = 0.001$ |
| **Relational accuracy** | $r = 0.147^*$<br>$p = 0.040$ | $r = 0.259^{**}$<br>$p < 0.001$ | $r = 0.218^{**}$<br>$p = 0.002$ | $r = 0.215^{**}$<br>$p = 0.003$ |
| **Progressive Matrices** | $r = 0.128$<br>$p = 0.075$ | $r = 0.327^{**}$<br>$p < 0.001$ | $r = 0.231^{**}$<br>$p = 0.001$ | $r = 0.271^{**}$<br>$p < 0.001$ |
| **Composite Cognitive Control Metric** | $r = 0.182^*$<br>$p = 0.011$ | $r = 0.442^{**}$<br>$p < 0.001$ | $r = 0.224^{**}$<br>$p = 0.002$ | $r = 0.277^{**}$<br>$p < 0.001$ |
| **Cohort 2 – IC3** | **Cortical thickness** | **Resting state** | **Relational tfMRI** | **2-Back tfMRI** |
| **Flanker task** | $r = 0.142$<br>$p = 0.084$ | $r = 0.122$<br>$p = 0.139$ | $r = 0.171^*$<br>$p = 0.037$ | $r = 0.216^{**}$<br>$p = 0.008$ |
| **2-Back accuracy** | $r = 0.098$<br>$p = 0.235$ | $r = 0.116$<br>$p = 0.158$ | $r = 0.248^{**}$<br>$p = 0.002$ | $r = 0.318^{**}$<br>$p < 0.001$ |
| **Relational accuracy** | $r = 0.211^{**}$<br>$p = 0.010$ | $r = 0.154$<br>$p = 0.062$ | $r = 0.373^{**}$<br>$p < 0.001$ | $r = 0.364^{**}$<br>$p < 0.001$ |
| **Progressive Matrices** | $r = 0.256^{**}$<br>$p = 0.002$ | $r = 0.260^{**}$<br>$p = 0.001$ | $r = 0.341^{**}$<br>$p < 0.001$ | $r = 0.317^{**}$<br>$p < 0.001$ |
| **Composite Cognitive Control Metric** | $r = 0.254^*$<br>$p = 0.002$ | $r = 0.234^{**}$<br>$p = 0.004$ | $r = 0.408^{**}$<br>$p < 0.001$ | $r = 0.437^{**}$<br>$p < 0.001$ |

### Replication by prediction: cross-cohort IC application

In order to further examine the replicability of these findings, the set of ICs from Cohort 1 (fig. 1B) were applied to Cohort 2’s source data (fig. 1D, left) and used to generate a set of subject specific weights for Cohort 2 (fig. 1E). These weights corresponded to the extent to which the data from a given subject in Cohort 2’s could be represented by the components from Cohort 1. These derived-weights replicated the significant correlation results identified in Cohort 1’s IC2 for three of the four modalities even after FDR correction (derived results for sMRI were not significantly correlated) (fig. 3 green data, table 3). That is, ICs containing measures of rsfcMRI, relational tfMRI, and two-back tfMRI data derived from Cohort 1 which were then applied to imaging data in Cohort 2 were highly and significantly correlated with cognitive control behavioral performance in Cohort 2.

**Table 3.**
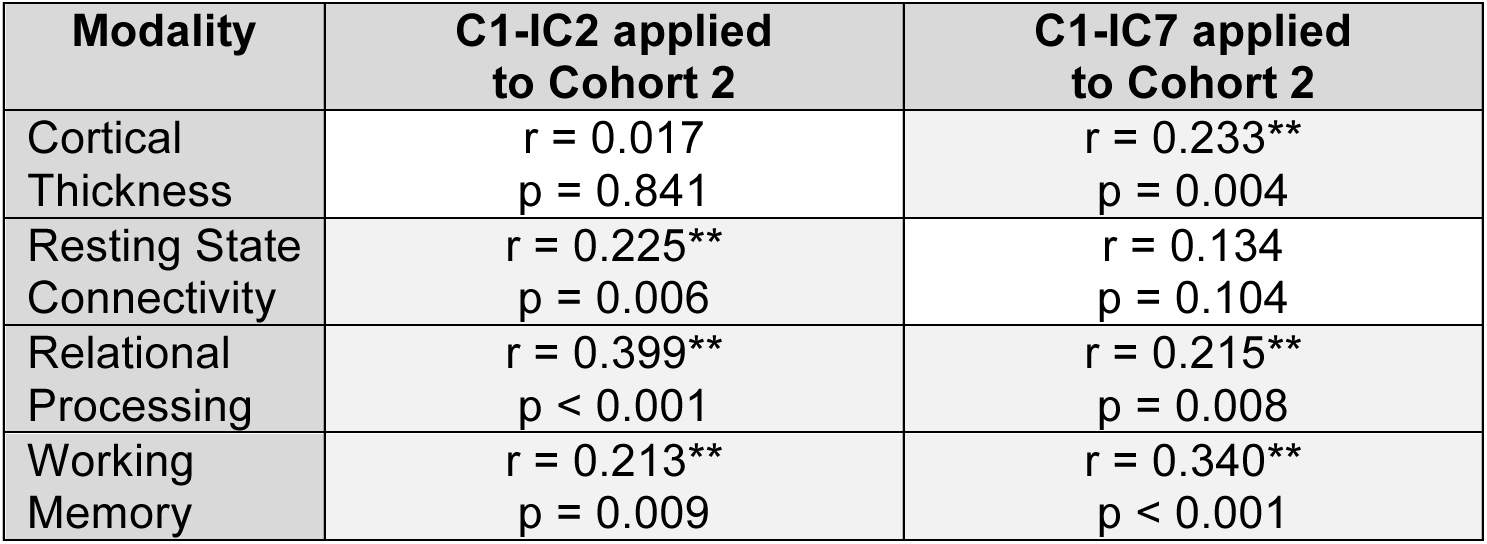
Correlations in Cohort 2 between cognitive control performance and subject specific weights derived from the replication by prediction analyses. <u>Figure Caption</u>: Correlations between the cognitive control composite measure and the subject-specific derived weights generated by using Cohort 1’s IC2 to decompose data from Cohort 2 replicate the significant correlations for three of the four modalities. FDR was computed using the p-values for all other correlations between the derived-weights and cognitive control (FDR critical p-value = 0.0264). Correlation results for the other ICs are in the supplement. *p < 0.05, **p < 0.01, uncorrected.

| Modality | C1-IC2 applied to Cohort 2 | C1-IC7 applied to Cohort 2 |
| --- | --- | --- |
| Cortical Thickness | $r = 0.017$<br>$p = 0.841$ | $r = 0.233^{**}$<br>$p = 0.004$ |
| Resting State Connectivity | $r = 0.225^{**}$<br>$p = 0.006$ | $r = 0.134$<br>$p = 0.104$ |
| Relational Processing | $r = 0.399^{**}$<br>$p < 0.001$ | $r = 0.215^{**}$<br>$p = 0.008$ |
| Working Memory | $r = 0.213^{**}$<br>$p = 0.009$ | $r = 0.340^{**}$<br>$p < 0.001$ |

### Replication by prediction: identification of a second IC correlated with behavior

Cohort 1’s independent component 7 (C1-IC7) was significantly correlated with behavior for the same three of the four modalities across both cohorts in the replication by prediction analysis (rsfcMRI data were not significantly correlated). No other component was significantly correlated with behavior across cohorts for the same three out of four imaging modalities. As shown in figure 4, sMRI data showed strong positive and negative contributions in a diffuse distribution across many cortical areas, with the strongest positive contributions observed in the bilateral insula and negative contributions in the left posterior and middle cingulate. Both relational and two-back tfMRI data showed broad regions of positively contributing vertices in bilateral medial and lateral prefrontal, inferiorparietal, insular, precuneus, and right middle temporal cortex. Both tfMRI modalities also shared negatively contributing vertices in bilateral visual, left precentral, and right supramarginal cortex. Relational data had additional positive contributing vertices in the left postcentral and inferior temporal cortex and negative contributing vertices in the right cuneus. Two-back data had additional negative contributing vertices in the left medial superior frontal and right posterior cingulate cortex.

### Replication by independent analysis

We also wanted to determine whether the independent application of mCCA+jICA to Cohort 2’s imaging data would replicate the findings from Cohort 1. In line with the parameter sweep methodology (see *supplementary methods*), the data from Cohort 2 were decomposed into nine ICs. Given the stochasticity inherent to mCCA+jICA, we algorithmically matched ICs across the two cohorts using the magnitude of the values within the ICs using eta^2^ (table S3). Of all nine matched pairs, the most similar pairing across the two cohorts was between Cohort 1’s second IC (C1-IC2) (described above) and Cohort 2’s third IC (C2-IC3), with 82.5% of variance explained. We examined the remaining ICs in Cohort 2, however none were a good visual match with Cohort 1’s IC7 (see figs. S13-S14 for the two closest matching ICs in Cohort 2). The eta^2^ values of the other 80 possible pairings are shown in table S3. Additionally, the same eta^2^ computation between cohorts was also performed for each modality independently. For all modalities, the similarity maximization algorithm again matched C1-IC2 with C2-IC3 (eta^2^ values: sMRI=90%; rsfcMRI=67%; relational tfMRI=80%; 2-back tfMRI=73%). Further, for sMRI, relational tfMRI, and 2-back tfMRI, the matching between C1-IC2 and C2-IC3 had the highest value of eta^2^ compared to all other matched pairs within the modality (rsfcMRI C1-IC2 and C2-IC3 had the second highest value of eta^2^). This suggested that the matching between C1-IC2 and C2-IC3 was driven by data from all four modalities rather than a subset of modalities. Further, the subject-specific imaging weights on all four modalities in C2-IC3 were significantly correlated with cognitive control performance (tables 1, 2).

In line with the eta^2^ results, visual examination of C2-IC3 revealed highly similar patterns to those observed in C1-IC2 (fig. 2, right column). C2-IC3 was visualized using the same |Z| > 2 threshold (Sui, Pearlson et al. 2015), and overlap maps were generated to aid visual comparison (figs. S6-S12). sMRI data for Cohort 2 were fully positive (as was the case with Cohort 1), and thus only values exceeding Z > +2 are shown. For sMRI, both cohorts had contributing areas in medial superior frontal cortex and superior and inferior temporal gyri, but the exact spatial locations did not overlap. For relational tfMRI data, Cohort 2 exhibited less positive contributions from the bilateral superior precentral sulcus and gyrus, and inferiotemporal and fusiform gyri. While both cohorts exhibited positive contributing areas in bilateral rostral middle frontal sulcus, the clusters were centered at slightly different locations. For 2-back tfMRI data, Cohort 2 exhibited additional positive contributing areas in the bilateral rostral middle frontal gyrus and bilateral superior parietal cortex. Data at a threshold of |Z| > 1 in the subcortex and cerebellum are available for reference in the supplement (figs S11-S12) (see *Limitations*). rsfcMRI data for Cohort 2 showed similar patterns of contributing connections to Cohort 1 with slightly greater extent of contributions than Cohort 1 from within-network connectivity in the DMN, cingulo-opercular, and dorsal attention networks.

## Discussion

The goal of the present study was to perform a data-driven analysis and replication of the multimodal neural correlates of cognitive control in a healthy community sample. We identified an independent component from the imaging data in Cohort 1 (C1-IC2) that was significantly correlated with cognitive control performance across all four imaging modalities used in this study. Further, these results replicated to a second cohort of subjects using two methods. When the imaging results for C1-IC2 were applied to the second cohort (replication by prediction), three of the four imaging modalities were also significantly correlated with cognitive control performance in the second cohort (measures of cortical thickness did not significantly correlate). An independent analysis of the second cohort identified a component, C2-IC3, which was highly similar to C1-IC2 and also significantly correlated with cognitive control performance for all four imaging modalities. Furthermore, the replication by prediction analysis identified a second component in Cohort 1, C1-IC7, which was significantly correlated with cognitive control performance for three of the four imaging modalities (cortical thickness did not significantly correlate) and, when applied to Cohort 2, was significantly correlated with cognitive control performance for the same three imaging modalities. However, an analogous component was not identified in the independent analysis of Cohort 2.

As mentioned in the introduction, much work has been performed to identify the neural underpinnings of cognitive control, primarily through single modality studies. This work has frequently implicated prefrontal cortical structures as the regions supporting cognitive control in all modalities used in this study. In fact, in their seminal paper on prefrontal cortex (PFC) involvement in cognitive control, (Miller and Cohen 2001) posit that structures within the PFC support cognitive control through the generation of bias signals that project to other neural systems and, in doing so, shift those processing areas to achieve the desired task outcome. Later work by Dosenbach and colleagues (Dosenbach, Visscher et al. 2006, Dosenbach, Fair et al. 2007) extended this model to include areas outside of PFC and subdivided cognitive control functionality into two systems, termed the fronto-parietal (FP) and cingulo-opercular (CO) networks. Under this model, the FP network functions on a shorter timescale and is responsible for initiation of task control and rapid updating in response to task demands and error signals. In contrast, the CO network functions on a longer timescale and is responsible for stable maintenance and updating of task rules and goals across trials (Dosenbach, Fair et al. 2007).

This model of cognitive control can be used a frame for interpreting the spatial distribution of C1-IC2 and C2-IC3 (fig. 2). The Miller and Cohen model suggests that signals from PFC function to bias processing regions. Interestingly, however, the tfMRI data in this component appears to capture relatively little PFC functionality. Outside of the PFC, it did capture positive contributions in the intraparietal sulcus involved in the fronto-parietal and dorsal attention networks (Gordon, Laumann et al. 2016), both of which are thought to be involved in rapid task control (Dosenbach, Fair et al. 2007). Instead of PFC contributions, both tfMRI modalities in C1-IC2 and C2-IC3 showed the strongest contributions from striate and extrastriate visual areas, which may be related to the visual processing demands of both tasks. Indeed, the two-back task required processing of multiple stimulus types including faces, places, tools, and body parts and the relational processing task involved comparisons across six shapes filled with six possible textures. Previous analyses in these data exhibited strong positive group-level activation in visual areas (Barch, Burgess et al. 2013), a finding that was replicated in the spatial maps of this component. The tfMRI data in these ICs were somewhat consistent with the whole-brain tfMRI meta-analysis by (Niendam, Laird et al. 2012), however this set of ICs identified greater contributions from visual areas and fewer from frontal areas. Again, this may be due to the highly visual nature of the two tfMRI modalities used in the present study. Indeed, Niendam and colleague’s work spanned 193 studies using a much wider variety of tasks than the present work.

The meta-analysis of structural correlates of PFC described in the introduction (Yuan and Raz 2014) identified an overall positive relationship between cortical thickness in the PFC and cognitive control performance. In contrast, examination of the sMRI data for C1-IC2 and C2-IC3 (fig. 2) showed strong positively contributing vertices in the anterior insula. The literature has assigned this region to several networks including the cingulo-opercular and salience networks (Dosenbach, Visscher et al. 2006, Dosenbach, Fair et al. 2007, Power, Cohen et al. 2011, Gordon, Laumann et al. 2016). Under the model presented in (Dosenbach, Fair et al. 2007), the anterior insula serves as a general "task-mode” controller functioning on a longer time scale and is responsible for integrating thalamic and prefrontal signals. This finding, in conjunction with the tfMRI findings, again points to the insula as playing a potentially important role in cognitive control. Nonetheless, it was somewhat puzzling that we did not also see contributions from PFC in the sMRI findings. It is possible that the relatively restricted age range of our healthy sample reduced variance in PFC metrics, and that structural variability in the PFC may be more apparent in samples with a wider age range.

Resting state (rsfcMRI) data from C1-IC2 and C2-IC3 (fig. 2) also showed significant contributions from a number of networks previously associated with cognitive control, including the fronto-parietal, cingulo-opercular, dorsal attention, and default mode networks. Interestingly, the rsfcMRI results appear visually similar to canonical resting state networks. That is, the resting-state networks identified here exhibit high within-network connectivity (sometimes termed modularity) as well as highly anticorrelated connectivity between the default mode network and task positive networks including the cingulo-opercular and dorsal attention networks. Given the positive correlation between cognitive control performance and the subject-specific weights upon the group-level rsfcMRI correlation matrix, the data suggest that individuals whose resting state networks more closely match "canonical” resting states may have better cognitive control performance, though we cannot make claims as to the directionality of this association. Nonetheless, our results are consistent with (Schultz and Cole 2016) who found a positive correlation between relational processing task performance and the extent to which a given subject’s task functional connectivity networks were similar to group-level resting-state functional connectivity networks. While these two results do not address the exact same question, together they suggest that there are behavioral performance benefits related to resting state network organization.

As noted, we also identified a second IC in Cohort 1, C1-IC7 (fig. 4), which was also correlated with cognitive control performance. This component was also correlated with Cohort 2’s cognitive control performance when the IC was directly applied to that cohort’s imaging data. However, an analogous component was not identified in the independent analysis of Cohort 2. In contrast to the predominantly posterior tfMRI contributions seen in C1-IC2 and C2-IC3 (fig. 2), C1-IC7 showed strong contributions distributed across lateral and medial dorsal aspects of the cortex. These findings were quite consistent with the regions identified in the cognitive control tfMRI meta-analysis by (Niendam, Laird et al. 2012). In both relational and 2-back tfMRI, there were strong positive contributions in the fronto-parietal and dorsal attention networks extending across dorsolateral and medial prefrontal cortex and posteriorly in the intraparietal sulcus (IPS) and portions of the inferior parietal lobule (IPL). Under the previously described models of cognitive control, these regions were associated with the fronto-parietal network and thought to be primarily reflective of the shorter timescale functionality of cognitive control. We identified further positive contributions in the precuneus and middle temporal lobe. While the Dosenbach model assigns the middle temporal lobe to a separate network, this region was seen in later models to associate with the fronto-parietal network (Power, Cohen et al. 2011, Thomas Yeo, Krienen et al. 2011, Gordon, Laumann et al. 2016). The involvement of the middle temporal lobe in this component, which contains predominantly rapid task-control network contributions, is consistent with the assignment of this region to the fronto-parietal network in later studies. Additionally, the Dosenbach model treats the precuneus as part of the fronto-parietal network which is consistent with our data, though later cortical parcellations treated it as a separate "parietal encoding and retrieval” network (Gordon, Laumann et al. 2016).

**Figure 4:**
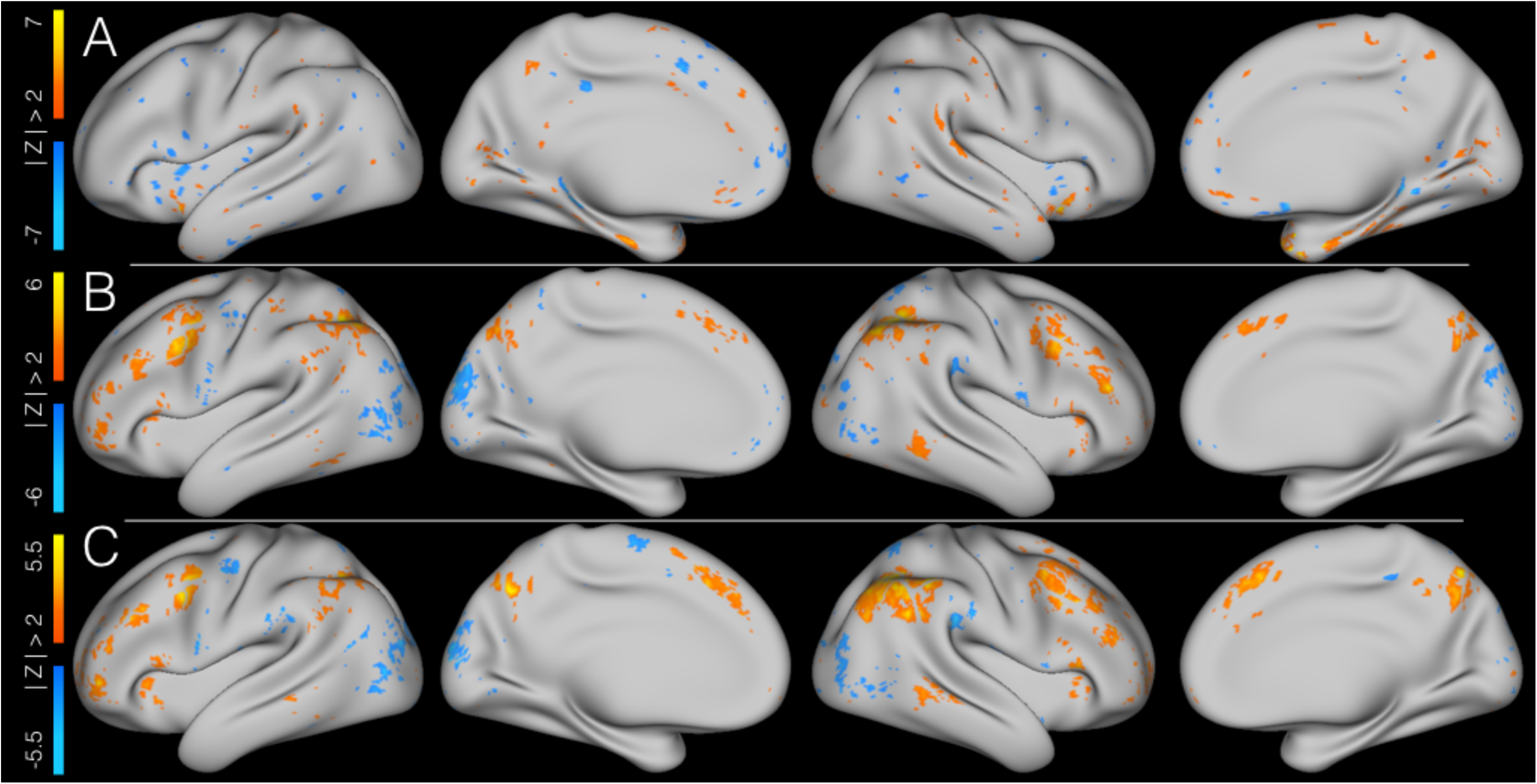
Imaging results for the modalities that significantly correlated with behavior in Cohort 1’s independent component 7. <u>Figure Caption</u>: Figure 4 shows the three modalities within Cohort 1’s independent component 7 that significantly correlated with cognitive control performance. All images show Z-scores of the IC spatial maps for a given modalities’ data and are thresholded at |Z| > 2. A = cortical thickness; B = relational tfMRI; C = 2-back tfMRI.

The sMRI contributions of C1-IC7 (fig. 4) were somewhat more difficult to interpret due to the relatively small clusters of contributing voxels and the somewhat scattered distribution across the cortex. The strongest positive clusters were located in portions of the anterior insula corresponding to the cingulo-opercular, salience, and ventral attention networks and strongest negative clusters were in the default mode and visual networks. Again, this finding is not consistent with the results of (Yuan and Raz 2014) in that we do not see strong contributions in PFC. However, it is interesting that both tfMRI modalities for C1-IC7, and the sMRI data for C1-IC7 and C1-IC2/C2-IC3 also showed contributions from the anterior insula. Under the Dosenbach model (Dosenbach, Fair et al. 2007), the anterior insula serves as a key hub in the cingulo-opercular network subserving the long timescale aspects of cognitive control. In our data, we consistently identified contributions from the anterior insula in components that were dominated by contributions from cognitive control networks associated with rapid timescale functions. One speculative hypothesis is that the involvement of the insula in both components may represent a role for the anterior insula in mediating between different networks and integrating the functions into a unified system. Indeed, previous data-driven meta-analytic analyses (Chang, Yarkoni et al. 2013) have suggested such a role for the insula.

As mentioned in the introduction, one of the open questions in the cognitive control literature is how findings in one modality relate to findings in other modalities. The results identified herein suggest that there may be a positive association between the various metrics of brain function and structure across modalities. That is, as identified in the similarity-matched IC pair, C1-IC2 and C2-IC3 (fig. 2), cortical thickness predominantly in the anterior insula, visually canonical resting state correlation matrices, positive task contributions from the visual and dorsal attention networks, and less task contributions from the default and cingulo-opercular networks were all jointly linked and these patterns all positively and significantly correlated with cognitive control performance (fig. 3). Similarly, for C1-IC7 (fig. 4), cognitive control performance was positively and significantly correlated with greater cortical thickness in the cingulo-opercular, salience, and ventral attention networks, less cortical thickness in the default mode and visual networks, strong task contributions in the fronto-parietal and dorsal attention networks, and less task contributions from visual networks (fig. 5). Thus, it is possible that these components, C1-IC2/C2-IC3 and C1-IC7, may reflect two major aspects of cognitive control: C1-IC7 may reflect contributions from regions that support dynamic aspects of task control and C1-IC2\C2-IC3 may reflect contributions from regions that support both stable and dynamic aspects of task control as well as regions that receive the influence of bias signals (i.e., more sensory regions).

**Figure 5:**
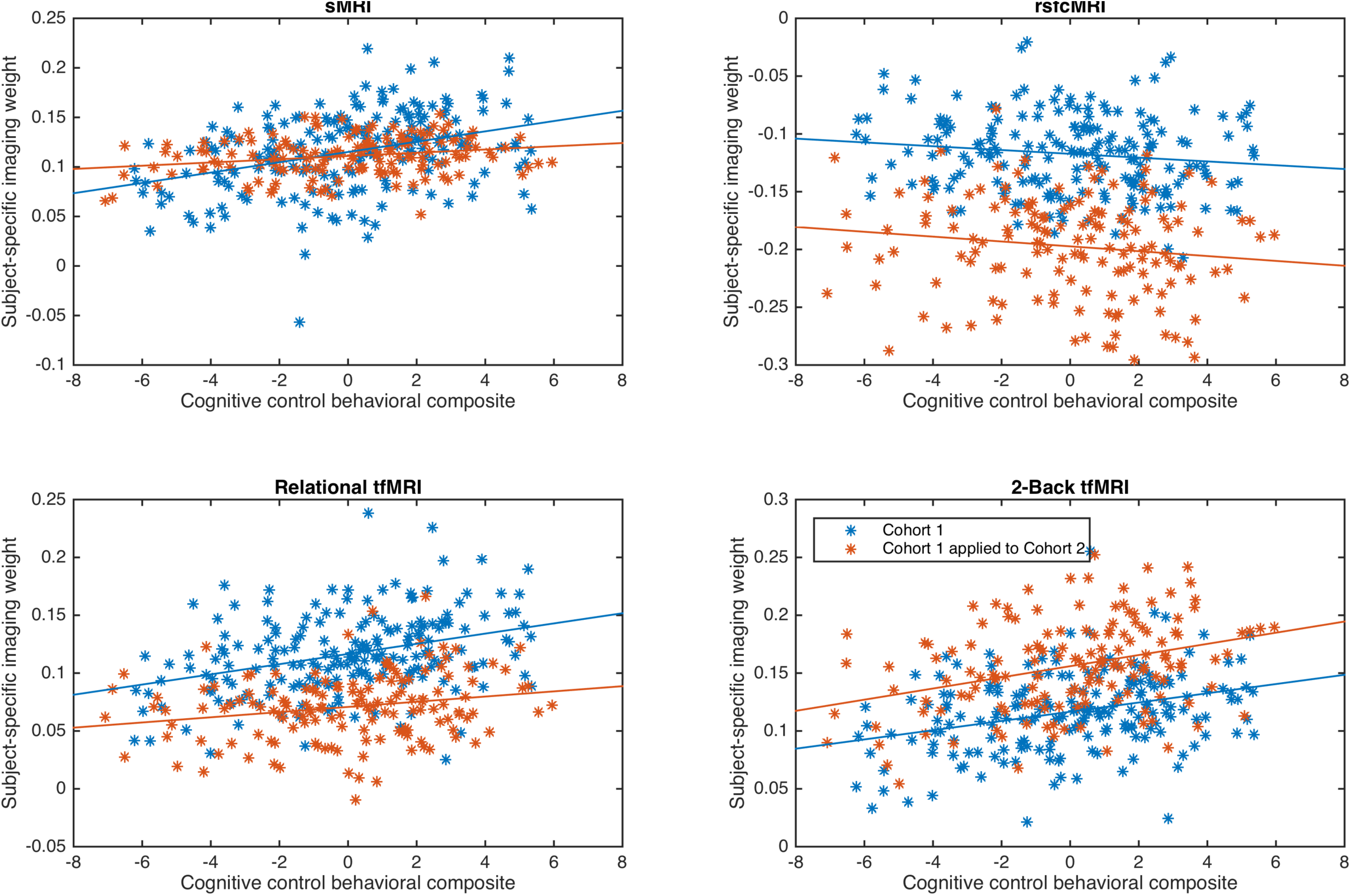
Scatter plots of cognitive control composite measure and subject-specific imaging weights for Cohort 1 independent component 7 and the replication by prediction analysis for Cohort 1 independent component 7 predicting Cohort 2. <u>Figure Caption</u>: Panels show the scatterplots and linear trendlines between the cognitive control behavioral composite and the subject-specific imaging weights on the respective imaging modality. Blue data are from Cohort1 and red from the application of Cohort 1’s IC7 to Cohort 2’s source imaging data. All correlations were statistically significant after FDR correction except for rsfcMRI data.

## Limitations and future directions

First, the present analyses were performed in a data-driven manner and caution should be taken in their interpretation and application to other datasets, though we were able to replicate our results in an independent cohort. Second, ICA-based methods are stochastic in nature and are unlikely to yield the exact same result when re-applied to a given dataset. However, the stability analyses we performed (see *supplemental methods* and *supplemental results*) to generate analysis model parameters suggests that the results presented herein are unlikely to be due to chance initial conditions in the decomposition. Additionally, the identification of highly similar components across the cohorts further suggests that the results are not due to chance initial conditions. While we did not identify an analogous component for C1-IC7, this may be due to the smaller number of subjects in cohort 2. Furthermore, while ICA-based methods are powerful tools for decomposing data, they do not guarantee perfect decomposition and separation of sources. This may explain why there was some inclusion of both fronto-parietal and cingulo-opercular results in the same components. Third, there was a relative paucity of strongly contributing voxels in the subcortex. This is likely due to the low signal-to-noise ratio in the subcortex arising from methodological choices in the collection of data for the human connectome project, and should be addressed by performing similar multimodal analyses in datasets with greater SNR in the subcortex. Future analyses should also include DTI data in order to assess the contributions of white matter and structural connectivity to cognitive control.

## Conclusion

The goal of the present study was to identify the multimodal neural correlates of cognitive control in a healthy community sample. We identified two imaging components in Cohort 1 that were highly correlated with cognitive control performance and partially replicated in a second independent cohort. The present findings were identified using data-driven methods to study the neural correlates of cognitive control and to help identify the relationships across modalities in a healthy community sample. Extending these findings to examine how these multimodal neural findings related to cognitive control are altered in psychopathology could yield key insights into the origins of deficits. Indeed, meta-analyses examining deficits in cognitive control have identified significant deficits in disorders such as attention-deficit/hyperactivity disorder (Willcutt, Doyle et al. 2005), antisocial behavior (Morgan and Lilienfeld 2000), major depressive disorder (Snyder 2013), and schizophrenia (Minzenberg, Laird et al. 2009). While studies of psychopathology have embraced neuroimaging as a tool to understand the source of behavioral manifestations, the predominance of studies have taken a unimodal approach and are thus unable to identify relationships in the data that may be present across modalities (Calhoun and Sui 2016). Such multimodal analyses in psychopathology could reveal whether deficits are more related to structural, functional, or connectivity alterations and thus provide novel targets for further research and intervention.

## Acknowledgements

We thank Malcolm Tobias, Keerthana Chivukula, and Anthony Chan for assistance with technical issues with MATLAB cluster computing. We also thank Jo Etzel for her excellent blog posts on using data from the Human Connectome Project <u>(http://mvpa.blogspot.com/)</u>.

